# The Voltage-gated sodium channel in Drosophila, Para, localizes to dendrites as well as axons in mechanosensitive chordotonal neurons

**DOI:** 10.1101/2023.03.07.531558

**Authors:** Thomas A. Ravenscroft, Ashleigh Jacobs, Mingxue Gu, Daniel F. Eberl, Hugo J. Bellen

**Author notes:** Corresponding Author – Hugo J. Bellen, DVM, PhD.

## Abstract

The fruit fly *Drosophila melanogaster* has provided important insights into how sensory information is transduced by Transient Receptor Potential (TRP) channels in the Peripheral Nervous System (PNS). However, TRP channels alone have not been able to completely model mechanosensitive transduction in mechanoreceptive chordotonal neurons (CN). Here we show that, in addition to TRP channels, the sole voltage-gated sodium channel (Na_V_) in *Drosophila*, Para, is localized to the dendrites of CNs. Para is localized to the distal tip of the dendrites in all CNs, from embryos to adults, and is colocalized with the mechanosensitive TRP channels No mechanoreceptor potential C (NompC) and Inactive/Nanchung (Iav/Nan). Para localization also demarcates spike initiation zones (SIZ) in axons and the dendritic localization of Para is indicative of a likely dendritic SIZ in fly CNs. Para is not present in the dendrites of other peripheral sensory neurons. In both multipolar and bipolar neurons in the PNS, Para is present in a proximal region of the axon, comparable to the axonal initial segment in vertebrates, 40-60μm from the soma in multipolar neurons and 20-40μm in bipolar neurons. Whole-cell reduction of *para* expression using RNAi in CNs of the adult Johnston’s organ severely affects sound-evoked potentials. However, the duality of Para localization in the CN dendrites and axons identifies a need to develop resources to study compartment-specific roles of proteins that will enable us to better understand Para’s role in mechanosensitive transduction.

**Significance Statement:** Several transient receptor potential (TRP) channels have been shown to localize to dendrites of *Drosophila* mechanosensitive chordotonal neurons (CN). Here, we show that the fly voltage-gated sodium channel, Para co-localizes with the TRP channels NompC and iav and a possible dendritic spike initiation zone (SIZ) in CN dendrites. This dendritic localization is unique to CNs, is not seen in other peripheral neurons, and may account for some aspects of mechanotransduction. Para also localizes to a SIZ at an axonal initial segment-like region, which is shared amongst many peripheral neurons.

## Introduction

Animals need to sense their environment to move, find food, and avoid predators. Peripheral nervous system (PNS) neurons are responsible for detecting environmental cues and relaying this information to the central nervous system (CNS). *Drosophila melanogaster* has provided important insights into sensory information processing in the PNS (Cosens and Manning, 1969; Göpfert et al., 2006; Akitake et al., 2015). The fly PNS contains multipolar and bipolar neurons (Bodmer and Jan, 1987; Orgogozo and Grueber, 2005). Multipolar neurons have one axonal neurite and many dendrites (Grueber et al., 2001; Wang et al., 2015). The expansive dendritic tree provides broad coverage of the animals’ periphery where transient receptor potential (TRP) channels open in response to pain, touch, and heat stimuli (Liu et al., 2003; Tracey et al., 2003; Zhong et al., 2010; Tsubouchi et al., 2012). Bipolar neurons have one axon and one dendrite separated by the soma (Orgogozo and Grueber, 2005). Bipolar neurons contain TRP channels sensitive to odors, chemicals, light, stretch, and sound (Stocker, 1994; Carlson et al., 1997; Vervoort et al., 1997; Ainsley et al., 2003; Schrader and Merritt, 2007; Suslak et al., 2015), and the singular dendrite enables the animal to precisely locate the direction of both attracting and deterring stimuli. Therefore, the TRP channel composition and orientation of neurons in the *Drosophila* PNS are optimized for sensing directional stimuli.

TRP channels alone do not account for all the electrophysiological properties of all sensory neuron dendrites. In *Drosophila,* a null allele for the mechanosensitive TRP channel *NompC*, which is localized to the most distal region of CN dendrites (Cheng et al., 2010; Lee et al., 2010), still has mechanosensitive properties in CNs (Eberl et al., 2000; Zhang et al., 2013). In addition to NompC, Drosophila CN dendrites also contain the TRP channels inactive (iav) and nanchung (nan) which form a functional heterodimer (Gong et al., 2004). Unlike NompC, nan and iav are required for the mechanosensitive response, however, in S2 cells the NAN-IAV complex alone is not mechanosensitive (Li et al 2021). Hence, it is likely that not all the ion channels responsible for mechanotransduction in *Drosophila* are known.

In another invertebrate, the crayfish *Astacus astacus*, a model of the mechanosensitive response in stretch receptors using just TRP channels was unable to recapitulate in vivo recordings (Swerup and Rydqvist, 1996). However, when the models were altered to incorporate Na_V_ channels, the recordings matched *in vivo* recordings indicating a possible mechanosensation role for Na_V_ channels (Suslak et al., 2011). Additionally, electrical spikes are present in mechanosensitive locust auditory neuron dendrites (Hill, 1983; Warren and Matheson, 2018) which are similar to the chordotonal neurons (CN) in the Johnston’s Organ (JO) in *Drosophila*. These spikes occur in the distal region of the dendrite and are sensitive to tetrodotoxin (TTX) suggesting a role for voltage-gated sodium channels (Na_V_) in mechanosensitive neurons (Hill, 1983; Warren and Matheson, 2018). Together this shows that mechanosensation in invertebrates does not just rely on mechanosensitive TRP channels and that Na_V_ channels may play a role in some peripheral neurons.

*Drosophila* has one Na_V_ channel gene encoded by *paralytic* (*para*) (Suzuki et al., 1971). In the unipolar neurons of the CNS, *para* is expressed in active, mature neurons and is localized to a SIZ at a distal axonal segment (DAS) (Ravenscroft et al., 2020). Little is known about Para distribution in the multipolar and bipolar neurons of the PNS. Gene expression reporters in late embryos reveal *para* is expressed in some PNS neurons but it is unclear if it is expressed in all or only some neurons (Hong and Ganetzky, 1994; Ravenscroft et al., 2020).

To determine the role of Na_V_ channels in mechanosensation in the *Drosophila* PNS we used a previously generated *Minos*-mediated integration cassette (MiMIC) protein trap inserted into the *para* locus to identify the distribution of Para (Venken et al., 2011; Lee et al., 2018; Ravenscroft et al., 2020). Using this allele, we identified the axonal SIZ of multidendritic and bipolar CNs in the 3^rd^ instar larval PNS in an axonal initial segment (AIS)-like region. Interestingly, we observe Para at the distal dendritic tip of all CN dendrites throughout development, indicating another role for Na_V_ channels in the peripheral mechanical response.

## Materials and Methods

### Fly lines and maintenance

Flies were raised on a standard molasses-based lab diet at 22°C in constant light conditions. All crosses were performed at 25°C in a 12-hour light/dark incubator. Animals were not selected for sex at embryonic, larval, or pupal stages. Fly lines used are listed in Table 1. All fly lines used are either deposited in Bloomington *Drosophila* Stock Center, the Vienna *Drosophila* Resource Center, or are available upon request. The characterization and validation of the gene-trap and protein-trap *para-alleles* were previously performed by Ravenscroft et al., 2020.

**Table 1.**
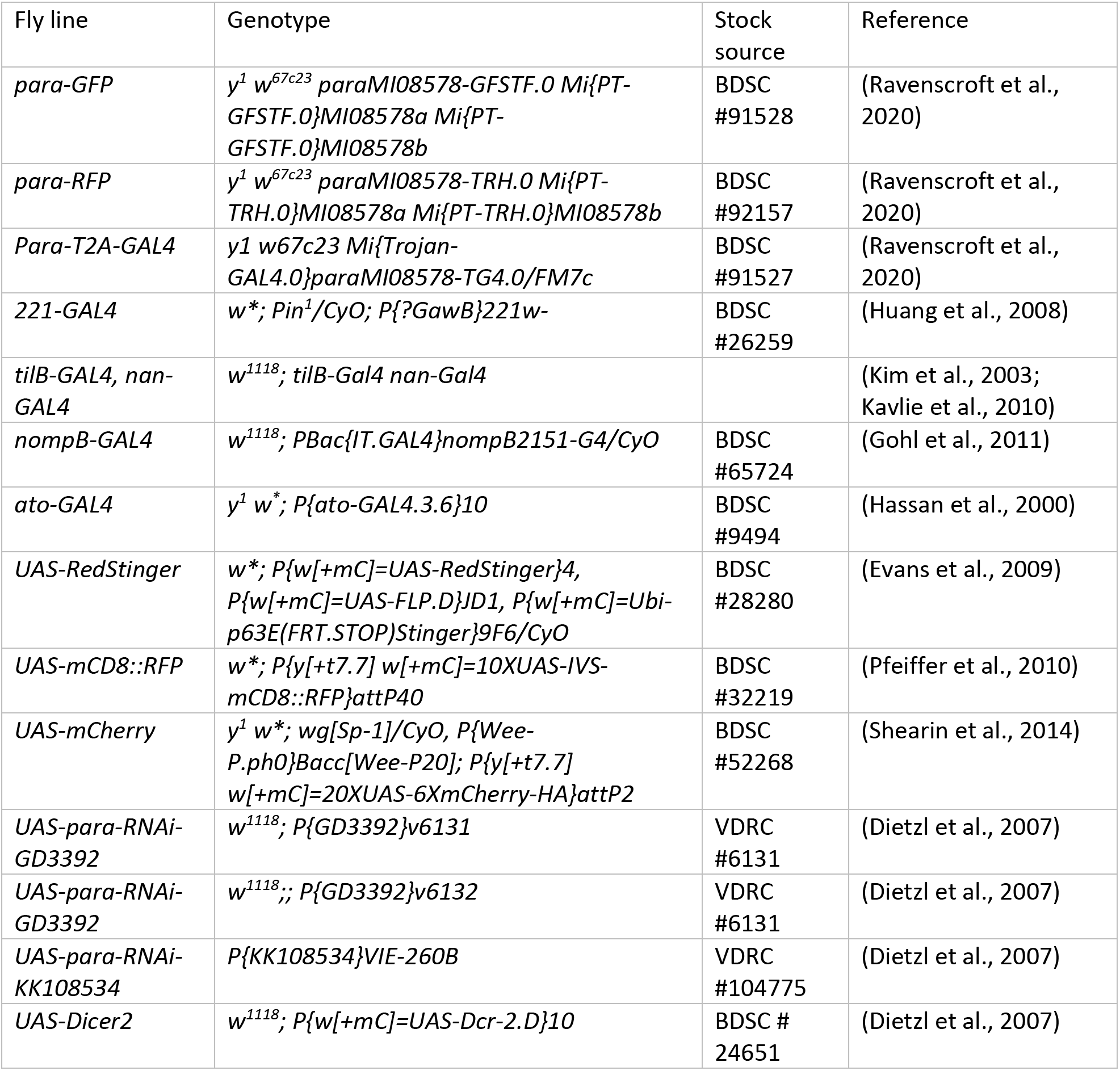
Summary of fly lines used in this study

### Immunostaining

#### Embryos

Immunostaining of Drosophila embryos was done as described previously (Rothwell and Sullivan, 2007). Flies were crossed in a chamber containing a grape juice plate (Welch) at 22°C in constant light. Flies lay eggs predominantly around dusk; therefore, to collect embryos at stage 16 we waited for 20-24 hrs. for collection. Embryos were collected with a paintbrush and water into a cell collection chamber (VWR #732-2758). These baskets were placed in a 50% bleach solution for dechorionation and agitated with a pipette. Dechorionation was observed under a microscope and once >75% of embryos lost the dorsal appendages the embryos were washed with an embryo wash solution (0.7%NaCl, 0.05% Triton X-100 in water). The basket was then dried by placing the chamber on a Kim wipe. For fixation, the mesh of the basket was removed with a razor blade and placed in a glass scintillation vial. The mesh was washed with 1ml of heptane saturated with 37% Formaldehyde (equal volumes of heptane and 37% formaldehyde were placed in a scintillation vial and vigorously mixed several times, the solution was allowed to settle into two phases with saturated heptane in the upper phase) which removed the embryos from the mesh. The mesh was then removed and 1ml of 3.7% formaldehyde in PEM buffer (0.1M PIPES, 1Mm MgCl2, 1mM EGTA, Ph 6.9, in water) was added, the vial was vigorously mixed for 15 s, and then left at room temperature for 20 mins. The bottom formaldehyde layer was then removed and replaced with methanol (100%), this mixture was vigorously mixed for 15 s and left to stand for 1 min. After the embryos sank to the bottom of the vial and the upper heptane layer was removed. The vial was then filled ∼2/3 full of 100% methanol and left to sit at 4°C overnight. Embryos were then transported to a 1.5 ml Eppendorf tube. As much methanol as possible was removed and replaced with 500 μl of PBTA (1X PBS, 0.05% Triton X-100, 0.02% Sodium Azide). They were then placed on a rotator at room temperature for 15 mins to rehydrate. Primary antibodies were then incubated in the vials in PBTA and left at 4°C overnight on a rotator. Antibodies were then recovered, and embryos were rinsed 3 times with PBTA and left for 1 hr. in PBTA on a rotator. Secondary antibodies were then added in PBTA and incubated on a rotator at room temperature for 2 hrs. (wrapped in foil to avoid light exposure). Antibodies were then recovered, and embryos were rinsed 3 times with PBTA and left for 1 hr. in PBTA on a rotator. Embryos were rinsed 4 times with PBS-Azide (1xPBS, 0.02% Sodium Azide) to remove the detergent. Embryos were mounted in rapiclear 1.47 (SUNjin labs) for imaging. Embryos were placed on a glass slide with a coverslip with no spacer. Antibodies used were rabbit-GFP 1:200 (Invitrogen #A-11122), rat-Elav 1:500 (DSHB #7E8A10), mouse-FLAG 1:200 (F3165 Sigma), rabbit-NompA 1:200 (Chung et al., 2001). Secondary antibodies used were goat-anti-HRP-Cy3 1:500 (Jackson ImmunoResearch Laboratories, #123-165-021) and corresponding donkey secondary antibodies 1:500 (Jackson ImmunoResearch Laboratories).

#### 3rd Instar Larvae

Wandering 3^rd^ instar larvae were placed in cold 1x Schneiders media (SM). Larvae were pinned on a Sylgard plate with Minuten pins in the anterior and posterior of the animal, dorsal side down. An incision was made with fine dissection scissors from the posterior to the anterior of the animal. Internal organs and fat were removed from the animal. Pins were placed to fillet the larvae. 3-4 animals at a time were filleted on each plate. The SM was then replaced with 3.7% PFA in SM and placed on a gentle rocker for 20 mins at room temperature. After 20 mins larvae were rinsed 3 times with SM, pins were removed and larvae were placed in a micro-Eppendorf tube in 0.1% PBS-TX (1X PBS, 0.1% Triton 100-X), tubes were washed 3 times for 10 mins on a rotator at room temperature. Primary antibodies were then added and incubated with 0.1% PBS-TX overnight on a rotator at 4°C. Antibodies were then recovered, and animals were rinsed 3 times with 0.1% PBS-TX and then washed 3 times for 10 mins in 0.1% PBS-TX. Secondary antibodies were then added and incubated (wrapped in foil) for 2 hrs. at room temperature on a rotator. Antibodies were then recovered, and animals were rinsed 3 times with 0.1% PBS-TX and then washed 3 times for 10 mins in 0.1% PBS-TX. Larvae were then mounted on a glass slide in rapiclear 1.47 (SUNjin labs) under a coverslip with no spacer. Antibodies used were rabbit-GFP 1:200 (Invitrogen #A-11122), mouse-GFP 1:200 (Sigma G6539), rabbit-NompA 1:500 (Chung et al., 2001), mouse-Eys 1:50 (DSHB #21A6 (Fujita et al., 1982)), rabbit-NompC 1:300 (Cheng et al., 2010). Secondary antibodies used were goat-anti-HRP-Cy3 1:500 (Jackson ImmunoResearch Laboratories, #123-165-021) and corresponding donkey secondary antibodies 1:500 (Jackson ImmunoResearch Laboratories).

Intensity profiles for Para distribution were calculated using a previously published approach (Jegla et al., 2016). Stacked confocal images of ddaE neurons were processed in ImageJ, a line was measured from the soma along the axon as far as a single axon track could be followed. The intensity of UAS-mCherry and GFP staining was recorded using the measurement feature. Relative intensity was measured by dividing the measured value by the average of the lowest 20% of measurements. This value was then divided by the top 5% of values (after dividing by the lowest 20%) to give a relative intensity. This was performed on n=15 neurons from n=5 animals. Measurements from all animals were then combined, smoothed out to an average of 50 values, and plotted on a graph. GFP to mCherry ratio was measured by comparing each relative GFP measurement to the corresponding mCherry measurement. For CN neurons the measurements were started at the distal tip of the dendrite through the soma into the axon. These data were represented with the soma at 0 using the max intensity point of mCD8::mCherry as the soma. Due to variation in dendrite length between animals, a representative trace is shown.

#### Johnston’s Organ

Johnston’s organ dissections and imaging were performed as described previously (Li et al., 2016). Pupae 24-48h after puparium formation were placed on double-sided tape on a glass slide. Using forceps, the outer shell was removed, and the pupae were removed and placed in a Sylgard dish in SM. Using micro scissors, the head was removed. A pipette was used to provide suction and remove the fat and the brain from the head, leaving the antenna attached to the outer membrane. This membrane was fixed in 3.7% PFA in PBS for 20 mins, then rinsed 3 times and washed 3 times for 10 mins in 0.1% PBS-TX. Samples were incubated in conjugated antibodies in 0.1% PBS-TX and incubated overnight at 4°C. Antibodies were recovered and samples were rinsed 3 times and washed 3 times for 10 mins in 0.1% PBS-TX. Samples were mounted in Vectashield on a glass slide with 2 pieces of double-sided tape acting as a spacer to protect the antennae. Antibodies used were rabbit-GFP-488 1:200 (Invitrogen), and Goat-Phalloidin-Cy3 1:500 (Jackson ImmunoResearch Laboratories).

Antibodies used were rabbit-GFP-488 1:200 (Invitrogen) and Goat-Phalloidin-Cy3 1:500 (Jackson ImmunoResearch Laboratories).

### Johnston’s organ electrophysiology

Sound-evoked potential (SEP) recordings were performed with an electrolytically sharpened tungsten recording electrode inserted into the joint between antennal segments one and two, and a reference electrode inserted into the head cuticle near the posterior orbital bristle, in response to near-field playback of computer-generated pulse song (described in Eberl and Kernan, (2011). The signals were subtracted and amplified with a differential amplifier (DAM50, World Precision Instruments) and digitized at 10 kHz (USB-6001, National Instruments). Average response values were measured as the max-min values in an averaged trace from 10 consecutive presentations of the described protocol.

### Experimental design

All confocal images of embryos were correctly aged using the structure of the CNS labeled by HRP. JO images were taken from pupae 48-72 hours after puparium formation. Adult flies aged 2-7 days were used for JO electrophysiology. Images from >5 animals for each condition were obtained.

### Statistical Analysis

A One-way ANOVA with Brown-Forsythe correction for unequal standard deviations was used for the comparison of SEP in JO electrophysiology data. Graphs indicate, within each bar, the number of antennae tested for that genotype. Statistical significance depicted on graphs indicate Tukey’s post-hoc multiple comparisons test.

## Results

### *para* is expressed in multidendritic neurons and CNs in Drosophila embryos

*para* expression is first noted at stage 16 of embryonic development where it is expressed in some CNS neurons (Hong and Ganetzky, 1994; Ravenscroft et al., 2020). The *para-T2A-GAL4* allele, when paired with a fluorescent reporter (*UAS-mcD8:GFP*), labels the cells that express *para* (Ravenscroft et al., 2020). We used *para-T2A-GAL4* to determine the expression pattern of *para* in the PNS in stage 16 embryos. The PNS of embryos contains developing multipolar and bipolar neurons responsible for mechanosensation, proprioception, temperature, and touch (Orgogozo and Grueber, 2005). Like in the CNS (Ravenscroft et al., 2020), *para* is not expressed in all PNS neurons (Figure 1A). *para* is expressed in a restricted number of multipolar neurons in the distal cluster of PNS neurons (Figure 1A’ box b). Unlike the multipolar neurons, *para* is expressed in all the bipolar CNs (Figure 1A’ box c). Note that muscle cells in the embryo are also not labeled with *para, in* contrast to vertebrates where Na_V_ channels are needed for muscle contraction (George et al., 1991).

**Figure 1.**
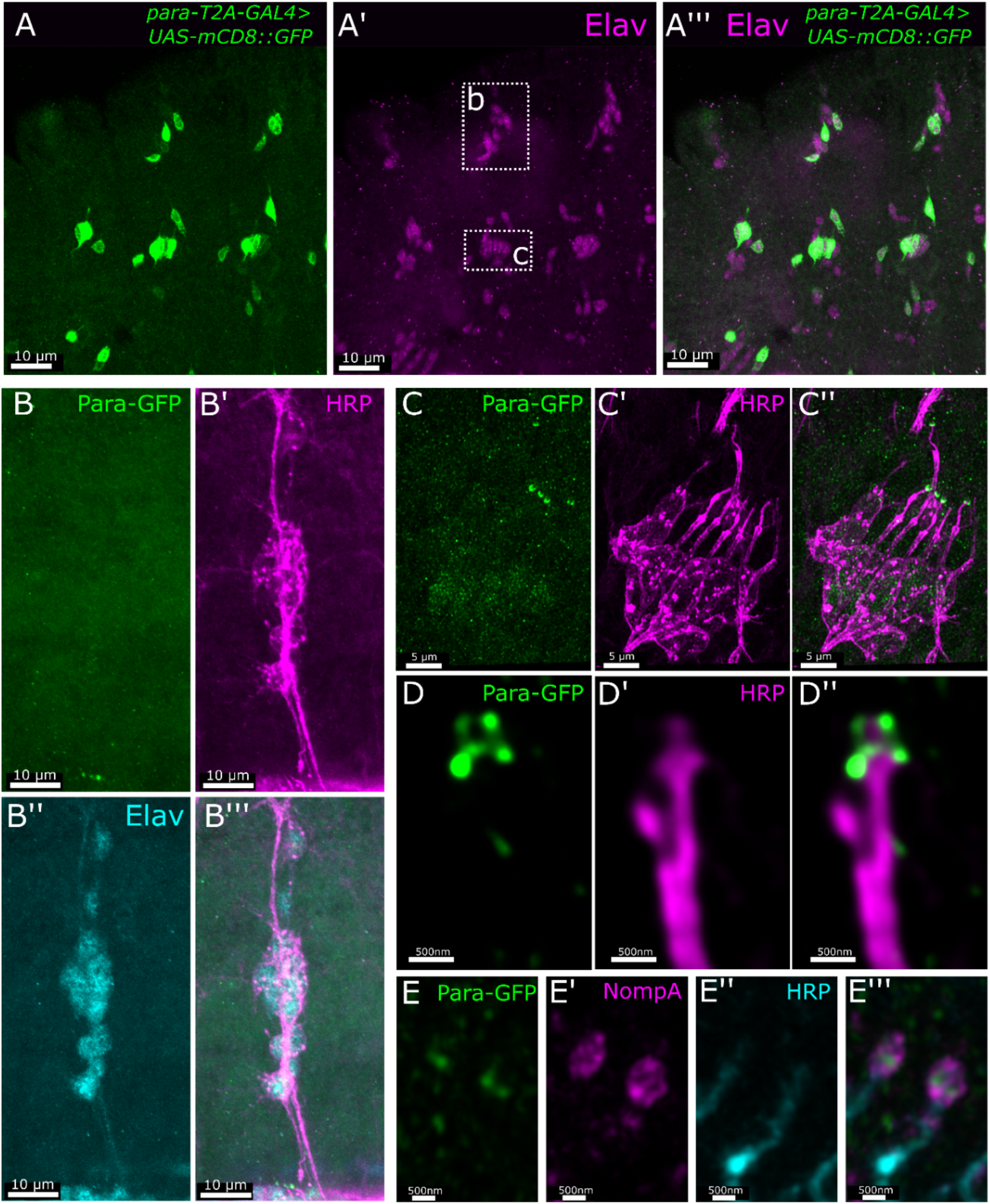
*para* is expressed in embryonic chordotonal neurons and is localized to dendrites. – the *para-T2A-GAL4* allele combined with a UAS-mCD8::GFP enables the visualization of *para-expressing* cells. (A) In stage 16 embryos, where *para* is first expressed, *para* is expressed in a restricted number of Elav-positive PNS neurons including all chordotonal neurons (box c). *para* is expressed in some multidendritic neurons (box b in A), despite this, the Para protein cannot be seen in axons, the soma, or dendrites indicating expression is very low (B). In the chordotonal neurons, Para is seen predominantly in the dendrites with less Para in the soma and no observed Para in axons. Neuronal membranes are labeled with an antibody against horse radish peroxidase (HRP) (C). Para is localized to the very distal tip of the dendrite (D), where it is surrounded by the dendritic cap protein NompA (E).

### Para is localized to embryonic CN dendrites and soma

To determine Para localization a MiMIC converted *para*-allele containing multiple epitopes for antibody labeling (Para-GFP-FlASH-Strep-TEV-FLAG (further referred to as Para-GFP)) was used (Venken et al., 2011; Ravenscroft et al., 2020). The Para-GFP allele has been previously validated and characterized to be representative of endogenous Para localization (Ravenscroft et al., 2020). We tested where Para is localized in the neurons relative to Elav which marks the soma of neurons and HRP which labels all neuron membranes.

In stage 16 embryonic multidendritic neurons, Para-GFP is not observed in the axons, dendrites, or soma (Figure 1B). At this stage the multidendritic neurons are still developing (Bodmer and Jan, 1987; Hartenstein, 1988); however, unlike the developing motor neuron of the CNS where Para is localized to the soma (Ravenscroft et al., 2020), Para is not detected in multidendritic neurons during development. This indicates Para’s functional role in these neurons occurs at later stages of development. CNs become fully differentiated in stage 16 of embryonic development (Jarman et al., 1993). *para* is strongly expressed in all embryonic CNs at stage 16. Para is predominantly localized to the distal tip of the dendrites with a lower level of Para seen in the soma (Figure 1C). Super-resolution stimulated emission depletion (STED) microscopy revealed that Para is present at the very distal tip of the dendrite with no membrane labeling via HRP present beyond where Para is localized in a basket-like structure ∼500nm from the dendrite tip (Figure 1D). No Para is detected in the axon at this stage. The distal tip of the CN dendrite is connected to a dendritic cap via an ECM that includes the glycoprotein NompA. NompA is specifically expressed in type I sense organs by support cells (scolopale cells) that ensheath the sensory process (Chung et al., 2001). In the distal CN dendrite of stage 16 embryos, Para is surrounded by NompA (Figure 1E).

### Para is expressed in all PNS neurons in 3^rd^ instar larvae and enriched at an AIS-like region in axons of multipolar neurons

We assessed the expression pattern in the 3^rd^ instar larval stage using *para-T2A-GAL4*. *para-T2A-GAL4* expression of *UAS-nls.mCherry* shows that *para* is expressed in all Elav-positive neurons in the PNS (Figure 2A). This contrasts with the expression of *para* in the 3^rd^ instar larvae CNS where it is only present in ∼25% of neurons (Ravenscroft et al., 2020).

**Figure 2.**
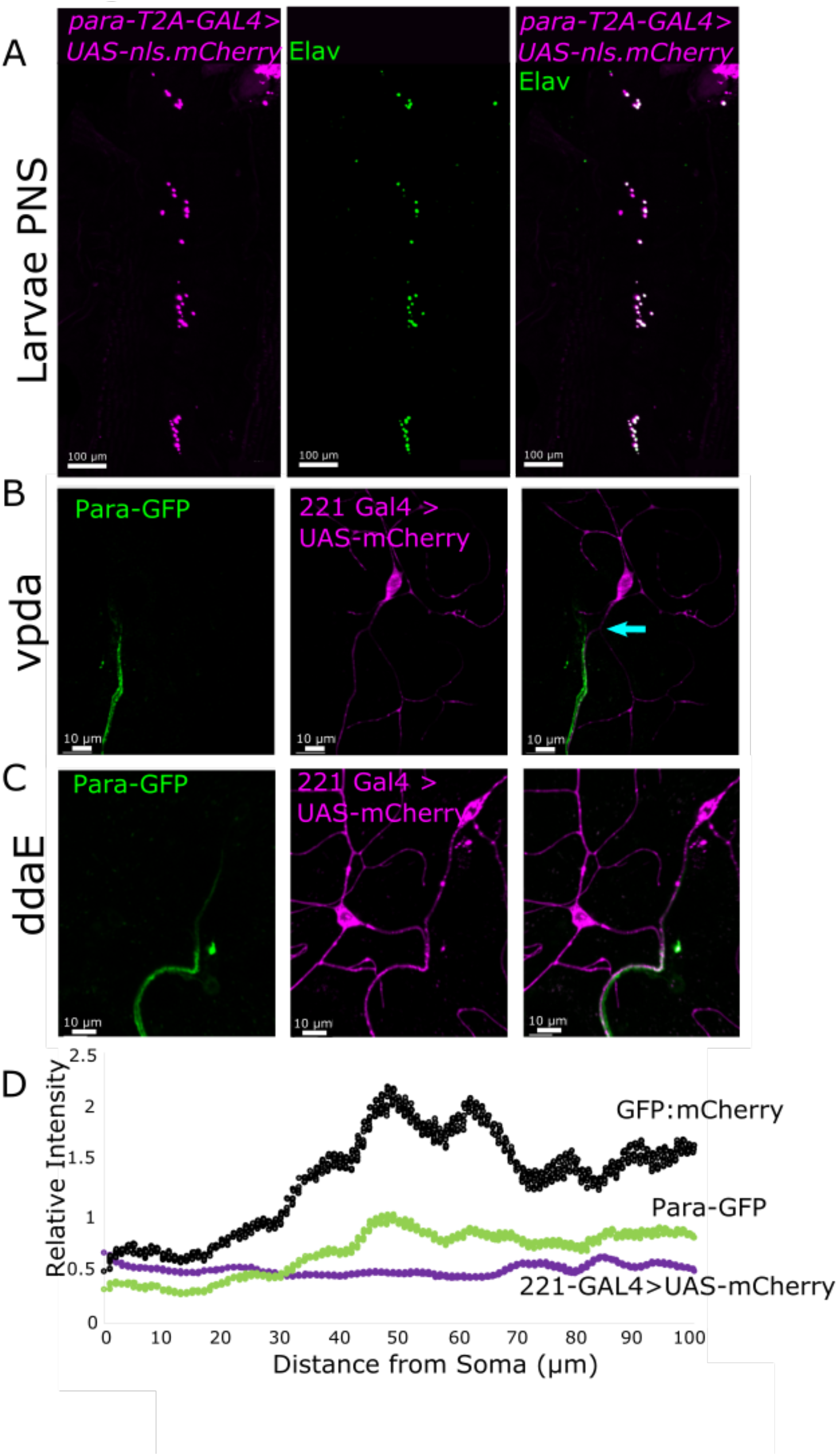
– Para is localized to an AIS-like region in multipolar neurons of the 3^rd^ instar larvae PNS. – *para* is essential for larval development and hence the expression and localization of Para in larval stages likely indicate where it functions. In the 3^rd^ instar larvae, PNS *para* is expressed in all neurons (Elav-positive cells) (A). In the 3^rd^ instar larvae multidendritic PNS neurons, Para is localized to the proximal axon in both vpda (B) and ddaE (C) neurons. In vpda neurons a dendrite can be seen in the proximal axon (arrow), Para is localized distal to this dendrite. In ddaE neurons, the relative intensity of Para localization is highest ∼ 40-60 μm from the soma indicating the presence of an axonal initial segment-like region in the 3^rd^ instar larva PNS (D). Beyond the AIS-like region, Para is still present at lower levels, likely to maintain the propagation of depolarizations to the synapse.

In the 3^rd^ instar larva and adult CNS Para is localized to a DAS while in stage 16 embryos Para is localized to the soma of neurons (Ravenscroft et al., 2020). Most CNS neurons in *Drosophila* are unipolar, while PNS neurons are either multipolar (multidendritic neurons) or bipolar (CNs). To determine Para localization in fully developed PNS neurons we used Para-GFP in combination with 221-GAL4, which drives GAL4 expression in some multipolar PNS neurons including vpda and ddaE neurons (Grueber et al., 2003), and UAS-mCherry in wandering 3^rd^ instar larva. In both the multidendritic vpda (Figure 2B) and ddaE neurons (Figure 2C), like CNS neurons, Para is localized to the axon but not the soma or dendrites. However, unlike in the CNS, Para is enriched in a segment that is only 40-60 μm from the soma (Figure 2D). This region overlaps with a previously reported AIS-like region marked by the localization of overexpressed Ank2 isoforms but is distal to the localization of overexpressed voltage-gated potassium (K_V_) channels Elk and Shal (Jegla et al., 2016). Beyond the AIS-like region, Para is still present at lower levels. We speculate that continued Para distribution is needed to maintain AP propagation beyond the initiation site. Additionally, in the vpda neuron, we observe dendrites that enter the axons beyond the cell body (Figure 2B). Similar to Para localization at a DAS in CNS neurons, Para is localized distal to the axonal dendrite in the vpda neurons.

### In the larva, PNS Para is localized to axons and dendrites of CNs

CNs are part of a four-cell chordotonal organ containing a neuron, a ligament cell that anchors the neurons, a cap cell that is attached to the CN dendrite via the dendritic cap, and a scolopale cell that protects and maintains the environment around the CN dendrite (Figure 3A)(Hartenstein, 1988). CN neurons and dendrites, like multidendritic neurons, continue to stretch and grow as the larvae grow (Singhania and Grueber, 2014). In wandering 3^rd^ instar larvae, the CNs reach their maximum length. In the lch5 CNs in the larval abdomen, Para is enriched in axons (Figure 3B). As in multidendritic neurons, Para in CNs is localized in the proximal part of the axon but unlike multidendritic neurons, Para is enriched close to the soma (20-30 μm) (Figure 3D). Additionally, the drop-off in the intensity of Para localization beyond the AIS-like region is greater in the CNs than in the multidendritic neurons. Hence, the AIS-like region is present in bipolar PNS cells as well as multipolar cells.

**Figure 3.**
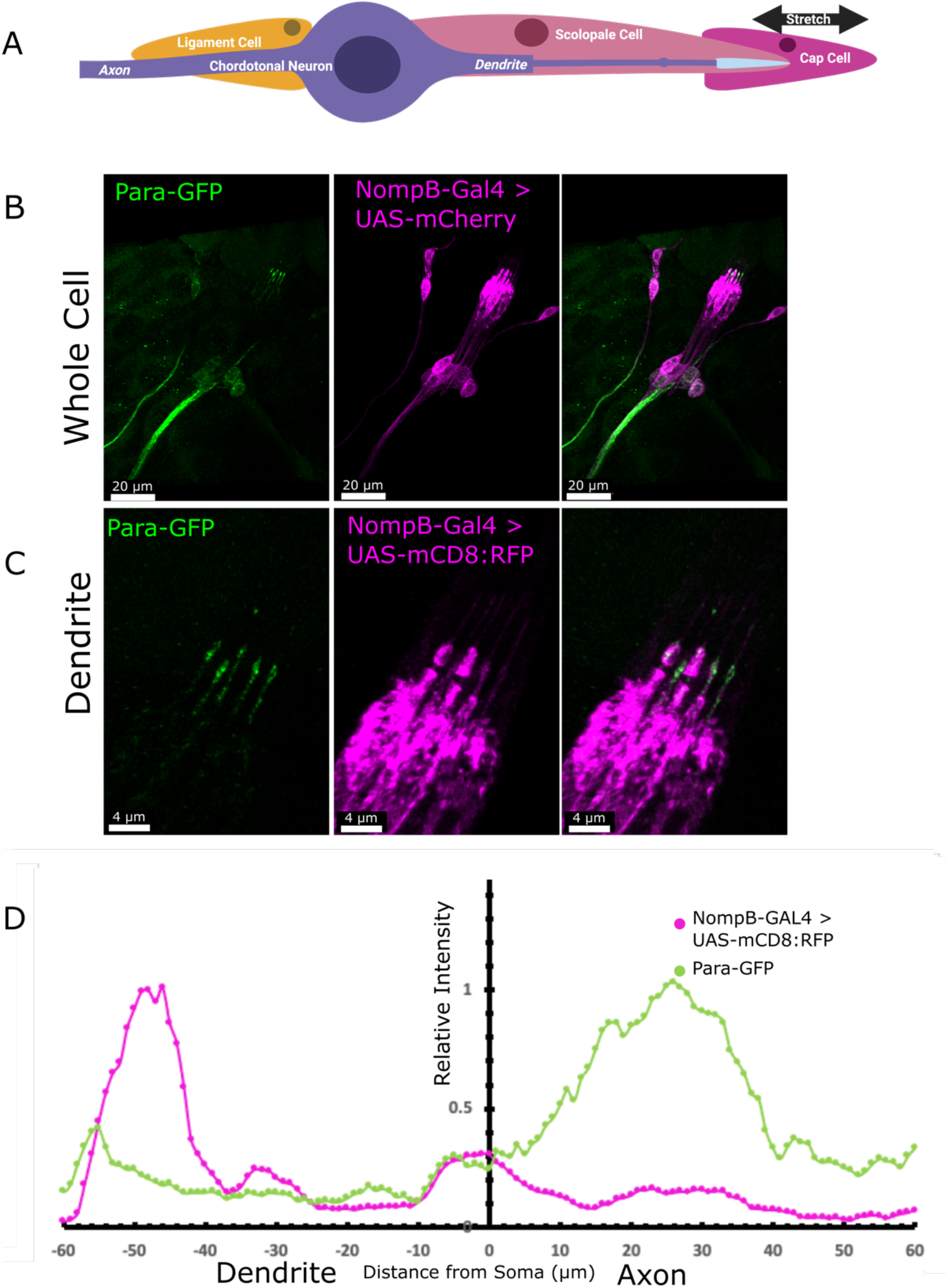
– Para is localized to both axons and dendrites in 3^rd^ instar larval chordotonal neurons. The chordotonal neuron is part of a 4-cell chordotonal organ (A). The neuron is anchored at the soma by a ligament cell and at the dendrite by the cap cell. The dendrite is insulated by a scolopale cell that provides structural support and protects the ionic balance of the dendritic space. NompB-GAL4 is expressed in the chordotonal neurons of the 3^rd^ instar larval PNS and with UAS-mCD8::RFP these neurons can be visualized (B). Para is enriched in both the axon and dendrite of the lch5 neuron (B). In the axon, Para is localized ∼20-30μm proximal to the soma, closer than what is observed in multidendritic neurons (D). In the dendrites (C), Para is localized to the distal dendrite ∼ 50-60μm from the soma (D). The distance on the axis in D indicates the distance from the soma into the dendrite (0μm to -80μm) and the axon (0μm to +80μm). 3A was created using biorender.com.

In the dendrites of the 3^rd^ instar larval lch5 CNs, like in embryonic CNs, Para is localized to the distal tip (Figure 3C). Para is less abundant at the dendrite than it is at the axon (Figure 3D). When compared to the localization of the cap protein NompA (Chung et al., 2001), Para is enriched both distal and proximal to the cap (Figure 4A). The more proximal localization of Para overlaps with the ciliary dilation which can be seen by the expansion of the dendrite distal to the extracellular scaffolding protein eyes-shut (Eys) (Figure 4B) (Blochlinger et al., 1991; Husain et al., 2006). Low levels of Para can also be seen proximal to the ciliary dilation. The dendrites of the CNs contain the mechanosensitive ion channels NompC, and the two interdependent TRP channels Inactive (Iav), and Nanchung (Nan) (Zhang et al., 2013). NompC is localized distal of the ciliary dilation while Iav and Nan are localized proximal to the ciliary dilation (Figure 4E) (Gong, 2004; Lee et al., 2010; Liang et al., 2011). Para predominantly colocalizes with NompC at the tip of the dendrite (Figure 4C). Para also partially colocalizes with the more proximal Iav (Figure 4D), however, the majority of Para in dendrites is in the distal region together with NompC.

**Figure 4.**
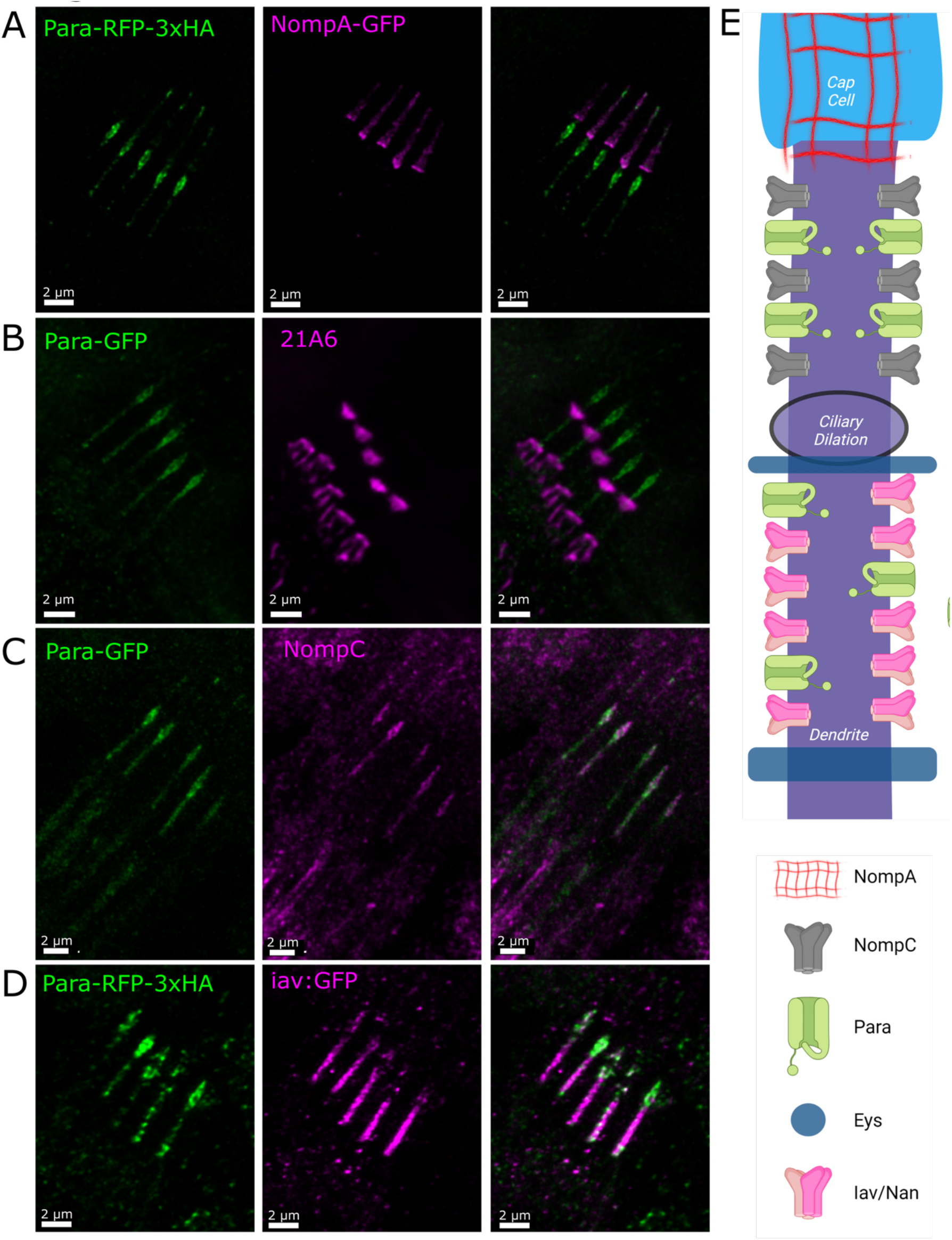
– Para colocalizes with NompC in the distal dendrite of 3^rd^ instar larva chordotonal neurons. Several proteins have been localized to dendrites of chordotonal neurons (E). NompA anchors the chordotonal dendrite to the dendritic cap. Para is localized to two segments of the dendrite, the most distal segment overlaps with NompA binding to the distal tip (A). The more proximal region of Para is on the proximal side of the dendrite where NompA is localized, distal to the scaffold protein Eyes-shut (labeled with Mab 21A6) (B). This location corresponds to ciliary dilation. The mechanosensitive transient receptor potential (TRP) channels NompC, Iav, and Nan are all localized to the dendrites of chordotonal neurons with NompC in the distal region and Iav and Nan proximal to the ciliary dilation. Para colocalizes with both NompC and Iav (C, D), however, Para is more abundant where NompC is localized in the distal dendrite. The localization of each protein is summarized in E. 4E was created using biorender.com.

### Para is required for sound response in the Johnston’s Organ CNs

Adult flies have a greater sensory repertoire than larvae. Adult flies have more sensitive responses to sound stimuli and have a greater abundance of proprioceptive and mechanosensitive neurons to maintain complex functions such as flight stabilization and courtship (Currier and Nagel, 2020; Montell, 2021). CNs are essential for hearing in adult *Drosophila* (Eberl et al., 2000). The 2^nd^ antennal segment of the adult fly contains the JO, which consists of 225 scolopidia (Caldwell and Eberl, 2002; Kamikouchi et al., 2006). Each scolopidium contains 2 or 3 bipolar CNs anchored in antennal segment 2 (Todi et al., 2004). The dendrites are attached to a tubular ECM dendritic cap anchored to the rotating stalk of antennal segment 3 (Todi et al., 2004). The neuron is surrounded by a scolopale cell, a glial-like cell that protects the dendrite and maintains the ionic balance of the extramembrane space of the dendrite (Caldwell and Eberl, 2002; Roy et al., 2013). When sound waves reach the antenna, they cause the stalk to rotate and pull on the dendrites of CNs, opening the mechanosensitive TRP channels NompC, Iav, and Nan and initiating a graded potential (Göpfert et al., 2006). Like the CN in larvae and embryos, *para* is expressed in all the CNs of the JO (Figure 5A). In these CNs, Para is also enriched in the dendrite at the distal tip and the ciliary dilation (Figure 5B, 5C).

**Figure 5.**
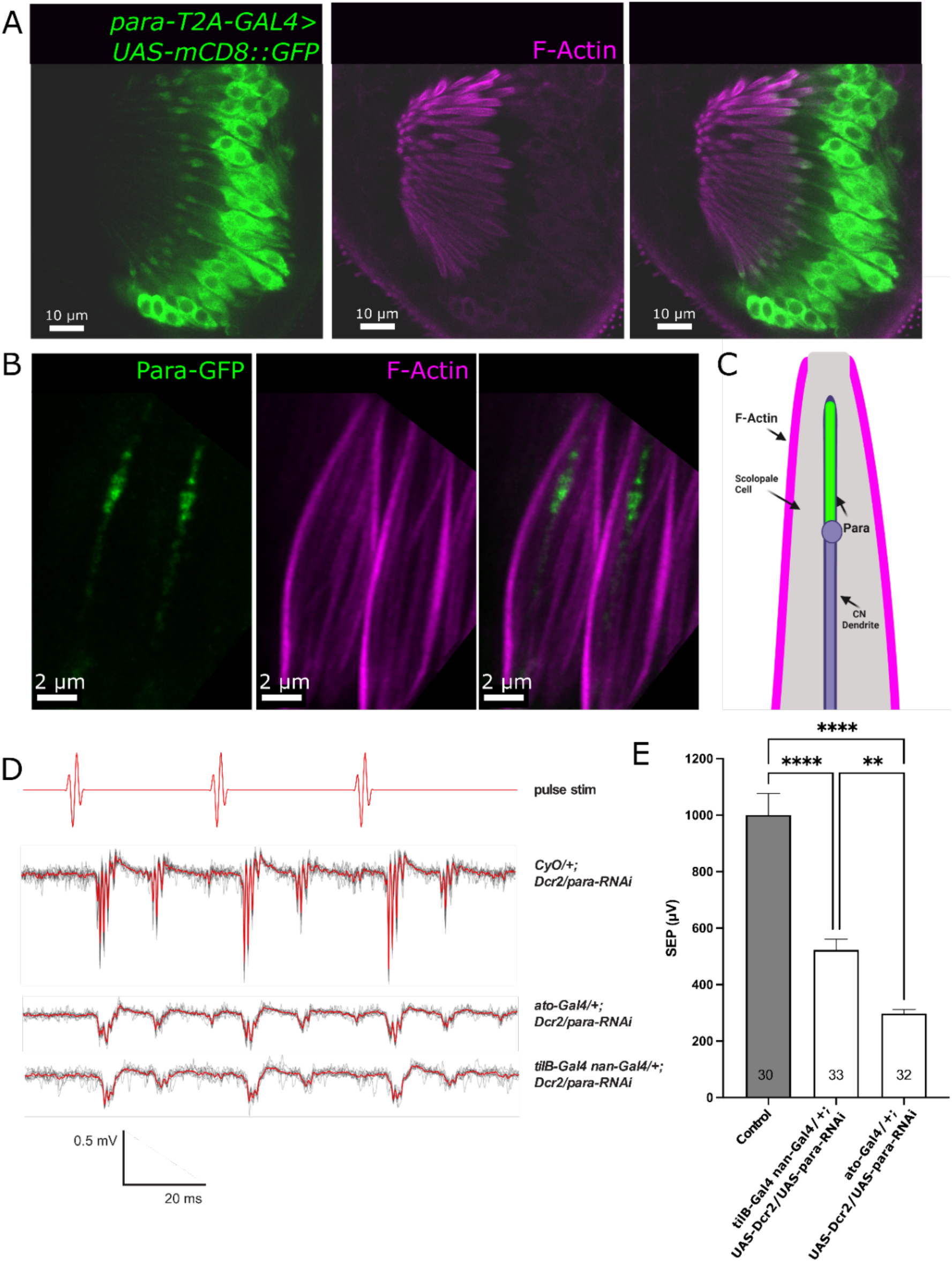
– Para is required for sound response in Johnston’s Organ (JO) - The JO is responsible for detecting auditory stimuli in the adult fly. In the developing pupae, ∼54 hrs after puparium formation *para* is expressed in the chordotonal neurons of the 2^nd^ antennal segment and not in the support scolopale cells labeled by F-Actin (A). In the chordotonal neurons, Para is localized to the distal dendrite of the chordotonal neurons in the JO (B). Diagram depicting the spatial arrangement of Para relative to the sensory cilium and scolopale rods (C). The activity of adult chordotonal neurons can be measured using sound-evoked potentials (SEP). Acoustic near-field presentation of computer-generated pulse song stimulus evokes strong SEPs in control animals with no Gal-4 driver. Responses to 10 individual stimulus presentations (depicted as light grey lines) are averaged (depicted as red lines) (D). RNAi knockdown of *para* using the *ato-Gal4* driver, which drives expression in the chordotonal sense organ precursor and the resulting chordotonal lineage, or with the *tilB*-Gal4 *nan*-Gal4 driver, which drives expression only in the chordotonal neurons, results in a strong reduction in the SEP amplitude (E). Bars and error bars indicate mean + SEM, and N’s shown within each bar indicate the number of antennae tested. Results of One-way ANOVA with Brown-Forsythe correction and Tukey’s post-hoc multiple comparisons test are shown (** p < 0.01; **** p < 0.0001). 5C was created using biorender.com. See also Supplemental Figure 5-1.

To determine if *para* is essential for CN mechanosensation in the adult JO, we used three RNAi lines against *para* driven by two separate CN drivers: *atonal-Gal4 (ato-GAL4),* which drives GAL4 expression in CN precursor cells and the chordotonal lineage, and *tilB-Gal4 nan-GAL4,* which drives GAL4 expression only in CNs (Kim et al., 2003; Kavlie et al., 2010; Roy et al., 2013). All RNAi lines target an exon incorporated into all 60 *para* isoforms ensuring we are not looking at isoform-specific effects (Larkin et al., 2021).

When *para* expression is reduced using any of the three RNAi lines against *para* with either GAL4 driver used, the sound-evoked potentials (SEP) produced in adult female flies in response to computer-generated male courtship pulse song is greatly reduced compared to control animals with no GAL4 driver (Figure 5D, 5E, Supplementary Figure 5-1). Additionally, when *para* expression is reduced, AP can no longer be detected in the SEPs (Figure 5D, 5E). Interestingly a small depolarization can still be detected in the neurons, possibly from the mechanosensitive channels remaining (NompC, Iav, Nan), however, when the *para* expression is reduced this signal is severely diminished implicating a role for Para in the excitability of the CN dendrite. Due to the dual nature of Para localization, the reduction in SEPs is likely the result of the loss of *para* in both axons and dendrites. However, we have not been able to isolate the dendrite or axonal-specific functions of para as we have not been able to remove Para just in dendrites. Attempts were made to inhibit Para using the sodium channel blocker tetrodotoxin, however, we were not able to access the CNs likely due to the glial sheath (Nelson and Laughon et al., 1993).

## Discussion

In *Drosophila*, the composition of ion channels that contribute to the graded potentials in PNS cell dendrites is unclear. Mapping the distribution of Na_V_ channels in the unipolar neurons of the fly CNS uncovered the SIZ at a DAS (Ravenscroft et al., 2020). Using the same endogenously tagged Para allele, we located the likely SIZ in the multipolar and bipolar neurons of the *Drosophila* PNS. In contrast to the DAS in the unipolar neurons of the fly CNS, the SIZ is at an AIS-like region proximal to the soma in PNS, comparable to the location of the AIS in vertebrate neurons (Huang and Rasband, 2018). Despite the more proximal location, the SIZ still determines the boundary between the somatodendritic and axonal compartments of the cell. Surprisingly, in addition to the axonal SIZ, a dendritic SIZ, demarcated by the presence of Na_V_ channels, is present in bipolar CN neurons. The dendritic SIZ is located at the distal tip of the dendrite and overlaps with the localization of the mechanosensitive TRP channel NompC and Iav. We believe this is a dendritic SIZ in agreement with a computational model of the crayfish CN for which Na_V_ activation is needed to accurately represent *in vivo* recordings (Suslak et al., 2011).

The localization of Para to the CN dendritic SIZ likely explains the TTX-sensitive dendritic spikes previously reported in insect mechanosensitive neurons (Hill, 1983; Oldfield and Hill, 1986; Lehnert et al., 2013; Warren and Matheson, 2018). Two TTX-sensitive spikes occur in the dendrites of locust auditory neurons dendrites, these spikes are recorded in the apical (distal) and basal (proximal) dendrite (Hill, 1983). The basal spikes respond to axonal depolarization and are likely backpropagating APs originating from Para channels opening at the SIZ in the axon. The apical spikes were of unknown origin but are likely to be spikes initiated by Para at the dendritic SIZ. While the dendritic SIZ identified in this study is in the fly and not the locust, Na_V_ localization is comparable between insect species as indicated by grasshopper Para and *Drosophila* Para having similar localization patterns (Wang et al., 2020).

All three identified TRP channels in the CN dendrites, NompC, Iav, and Nan, contribute to the mechanotransduction response. Direct patch clamp recordings of lch5 neurons identify a complete loss of mechanotransduction in the absence of Iav and Nan, while loss of NompC did not decrease the mechanotransduction response indicating that Iav and Nan are the essential channels (Li et al 2021). Patch clamp experiments did uncover that without NompC the adaptation time in CNs is a lot shorter (Li et al 2021), therefore the interplay between TRP channels in the dendrites is key for their proper function. The presence of Para in the same dendritic space as both NompC and Iav would enable Para to facilitate this interaction. It is worth noting however that these patch-clamp experiments were performed in conditions where Na_V_ was inhibited, therefore the electrophysiological interplay between Para and the TRP channels is still unknown.

TRP channel opening typically occurs in response to mechanical, temperature, chemical, or noxious stimuli (Clapham et al., 2001; Montell et al., 2002; Zheng, 2013; Montell, 2021). However, many TRP channels have been shown to open in response to voltage (Hofmann et al., 2003; Nilius et al., 2003; Voets et al., 2004; Matta and Ahern, 2007). The rat TRP channels TRPV1 and TRPM8 are hot and cold-responsive respectively (Dhaka et al., 2006), however, under specific conditions, they can open in response to voltage (Voets et al., 2004; Matta and Ahern, 2007). At room temperature and physiological pH, TRPV1 opens at around 0 mV with a half-activation voltage of around 150 mV (Matta and Ahern, 2007). These activation ranges are a lot harder to achieve (compared to the properties of Na_V_1.2 (activation ∼ -55 mV, half-activation voltage ∼-15 mV (Ogiwara et al., 2009)). However, for these temperature-sensitive channels when the channel is exposed to higher temperatures the structure of the channel changes, and the half-activation threshold is far lower, -50mV for TRPV1 at 42°C, which is readily achievable in a sensory neuron (Voets et al., 2004). It is not known if the fly TRP channels are voltage sensitive, however, the two closest homologs of the vertebrate voltage sensing TRP channel *TRPV1* are *nan* (DIOPT 7/16) and *iav* (5/16) (Hu et al., 2011). Interestingly, *iav* and *nan* are only expressed in the chordotonal PNS neuron dendrites where Para is localized (Kim et al., 2003; Gong, 2004). The possible voltage sensitivity of iav/nan in the same location as a Na_V_ channel suggests that Para activation could influence iav/nan activation or vice versa.

The dendrite of CNs is exposed to stretch forces opening mechanosensitive TRP channels. The presence of Para in the CN dendrite imposes the question as to whether Para may also be mechanosensitive. Two vertebrate Na_V_ channels Na_V_1.4 (*SCN4A*) and Na_V_1.5 (*SCN5A*) that are expressed in contracting muscle tissue have accelerated activation and inactivation kinetics when stretched in *Xenopus* oocytes (Shcherbatko et al., 1999; Tabarean et al., 1999; Morris and Juranka, 2007). While *in vivo* evidence for these channels’ mechanosensitivity is lacking, the G615E mutation in *SCN5A* which predominantly causes long-QT syndrome in patients has normal voltage-gating but aberrant mechanosensitivity indicating a role for Nav1.5 in mechanosensitivity (Strege et al., 2019). While *para* shares homology with *SCN4A* (DIOPT 9/16) and *SCN5A* (DIOPT 11/16) (Hu et al., 2011), *para* is not expressed in muscle cells, and evidence for neuronal Na_V_ channel mechanosensitivity is less obvious (Morris, 2011). *para* has 60 isoforms with different protein sequences and different dynamic properties that likely accommodate the requirements of different neurons (Lin et al., 2009, 2012). An isoform-by-isoform expression study is needed to determine if the CN *para* isoforms are closer in homology to *SCN5A* and *SCN4A* and thus more likely to be mechanosensitive.

In vertebrate neurons, voltage-gated ion channels can also contribute to graded potentials (Stuart et al., 1997; Golding and Spruston, 1998; Losonczy and Magee, 2006). In cultured hippocampal CA1 pyramidal neurons and brain slices of rat neocortical pyramidal neurons, spikes in membrane potential are observed in dendrites (Stuart et al., 1997; Losonczy and Magee, 2006; Sun et al., 2014). These dendritic spikes are not affected by the Ca_V_ channel blocker cadmium but are blocked by the Na_V_ channel blocker TTX, indicating that they are also generated by Na_V_ channels (Stuart et al., 1997). In CA1 pyramidal neurons, Na_V_1.6 is found in the dendrites, but it is 40 times less abundant than it is in axons (Lorincz and Nusser, 2010), while in the dendrites of *Drosophila* CNs, the max intensity of Para is roughly half that seen in the axon (Figure 3D). Hence, Na_V_-dependent dendritic spikes are not an insect-specific phenomenon.

The inner hair cells of humans have a comparable organization to that of the fly CNs (Boekhoff-Falk, 2005). Studies in flies have elucidated key genes and mechanisms of the auditory response in humans (Li et al., 2018). One Na_V_ channel in vertebrates, *SCN8A*, has been implicated in a mouse model of peripheral hearing loss (Mackenzie et al., 2009). Interestingly, dominant variants in *SCN8A* are often implicated with more global neurodevelopmental disorders (Trudeau, 2006; Veeramah et al., 2012). The specific hearing loss identified in the prior mouse studies indicates a specific role in hearing for the affected residue. Na_V_ localization to the CNs in flies opens the door for the use of *Drosophila* to study the impact of Na_V_ dysfunction on hearing loss.

While we propose a role for Na_V_ channels in CN dendrites, the presence of Para in the AIS-like region in addition to the distal dendrites makes the delineation of Para’s role in either region impossible with current tools. Reduction of *para* expression using RNAi in the adult JO neurons shows a strong reduction of SEP indicating an inability to process sound. However, we are reducing Para in both the axons and the dendrites and therefore we cannot distinguish if the loss of Para in dendrites prevents the dendritic depolarization from reaching the soma or if the axonal AP is lost due to a lack of Para in the AIS. To answer this question new tools are needed to selectively remove proteins in a compartment-specific way.

Para is localized to an AIS-like region in the axons of *Drosophila* PNS neurons. The region of Para localization overlaps with the previously reported AIS-like region in multipolar ddaE neurons, identified by an accumulation of over-expressed Ank2, Shal, and Elk and a diffusion barrier akin to the one seen at the vertebrate AIS (Jegla et al., 2016). The localization of Para and the AnkG homolog Ank2 in the AIS-like region of the PNS is of note as the AnkG binding motif in Na_V_ channels is not present in *para* indicating an alternative binding site and/or clustering mechanism in the fly PNS AIS-like region (Jegla et al., 2016).

In this study, we have identified the likely SIZ(s) in the multi- and bi-polar neurons of the fly PNS through the characterization of Na_V_ channel distribution. We have confirmed the presence of an axonal SIZ at an AIS-like region and surprisingly identified a likely dendritic SIZ in CNs throughout *Drosophila* development. The presence of Na_V_ channels in a dendritic and axonal SIZ in the fly PNS introduces an accessible system for further study into the role of Na_V_ channels in how animals sense their environment.

## Acknowledgments

DFE would like to acknowledge Jason Caldwell for preliminary RNAi experiments. We would like to thank Changsoo Kim for the iav-GFP flies and Yuh-Nung Jan for the NompC antibody. Figures 3A, 4E, and 5C were created using Biorender.com by TAR. TAR was supported by The Cullen Foundation. DFE and AJ were supported in part by NSF 2037828. AJ was supported in part by a University of Iowa Center for Research by Undergraduates Fellowship. HJB was supported by the Howard Hughes Medical Institute.

## Supplementary Figure 5-1

**Supplemental Figure 5-1-.**
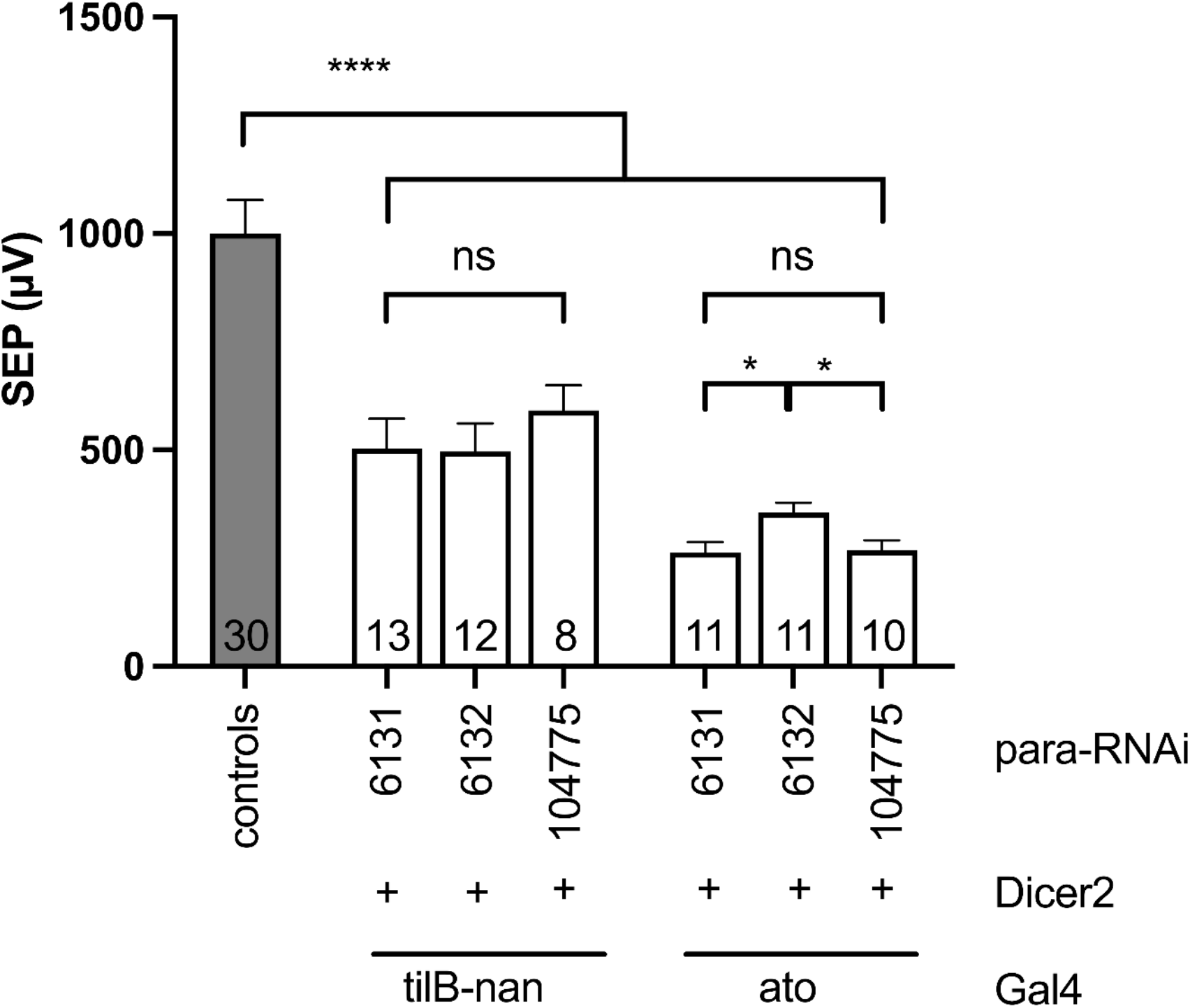
Multiple *para* RNAi lines reduce SEP when expressed in Johnston’s Organ chordotonal neurons. Electrophysiological tests as in Figure 5 show significantly reduced SEPs when driving 3 different RNAi lines (6131, 6132, 104775) using 2 different Gal4 drivers (a chromosome carrying both tilB-Gal4 and nan-Gal4, and ato-Gal4), all in the presence of UAS-Dicer2 (indicated by +). Symbols and statistics as in Figure 5. (ns, not significant; * p > 0.5; **** p < 0.0001).

## References

Ainsley JA, Pettus JM, Bosenko D, Gerstein CE, Zinkevich N, Anderson MG, Adams CM, Welsh MJ, Johnson WA (2003) Enhanced Locomotion Caused by Loss of the Drosophila DEG/ENaC Protein Pickpocket1. Current Biology 13:1557–1563

Akitake B, Ren Q, Boiko N, Ni J, Sokabe T, Stockand JD, Eaton BA, Montell C (2015) Coordination and fine motor control depend on Drosophila TRPγ. Nature Communications 6:7288

Blochlinger K, Jan LY, Jan YN (1991) Transformation of sensory organ identity by ectopic expression of Cut in Drosophila. Genes & Development 5:1124–1135

Bodmer R, Jan YN (1987) Morphological differentiation of the embryonic peripheral neurons in Drosophila. Roux’s Archives of Developmental Biology 196:69–77

Boekhoff-Falk G (2005) Hearing in Drosophila: development of Johnston’s organ and emerging parallels to vertebrate ear development. Dev Dyn 232:550–558

Caldwell JC, Eberl DF (2002) Towards a molecular understanding of Drosophila hearing. Journal of Neurobiology 53:172–189

Carlson SD, Hilgers SL, Juang JL (1997) Ultrastructure and blood-nerve barrier of chordotonal organs in the Drosophila embryo. Journal of Neurocytology 26:377–388

Cheng LE, Song W, Looger LL, Jan LY, Jan YN (2010) The Role of the TRP Channel NompC in Drosophila Larval and Adult Locomotion. Neuron 67:373–380

Chung YD, Zhu J, Han Y-G, Kernan MJ (2001) nompA Encodes a PNS-Specific, ZP Domain Protein Required to Connect Mechanosensory Dendrites to Sensory Structures. Neuron 29:415– 428

Clapham DE, Runnels LW, Strübing C (2001) The trp ion channel family. Nature Reviews Neuroscience 2:387–396

Cosens DJ, Manning A (1969) Abnormal Electroretinogram from a Drosophila Mutant. Nature 224:285–287

Currier TA, Nagel KI (2020) Multisensory control of navigation in the fruit fly. Current Opinion in Neurobiology 64:10–16

Dhaka A, Viswanath V, Patapoutian A (2006) TRP ion channels and temperature sensation. Annual Review of Neuroscience 29:135–161

Dietzl G, Chen D, Schnorrer F, Su K-C, Barinova Y, Fellner M, Gasser B, Kinsey K, Oppel S, Scheiblauer S, Couto A, Marra V, Keleman K, Dickson BJ (2007) A genome-wide transgenic RNAi library for conditional gene inactivation in Drosophila. Nature 448:151–156

Eberl DF, Hardy RW, Kernan MJ (2000) Genetically Similar Transduction Mechanisms for Touch and Hearing in *Drosophila*. The Journal of Neuroscience 20:5981–5988

Eberl DF, Kernan MJ (2011) Recording Sound-Evoked Potentials from the *Drosophila* Antennal Nerve: Figure 1. Cold Spring Harbor Protocols 2011:prot5576

Evans CJ, Olson JM, Ngo KT, Kim E, Lee NE, Kuoy E, Patananan AN, Sitz D, Tran P, Do M-T, Yackle K, Cespedes A, Hartenstein V, Call GB, Banerjee U (2009) G-TRACE: rapid Gal4-based cell lineage analysis in Drosophila. Nature Methods 6:603–605

Fujitya SC, Zipursky SL, Benzer S, Ferrús A, Shotwell SL (1982) Monoclonal antibodies against the Drosophila nervous system. Proceedings of the National Academy of Sciences 79(24)7929–33

George AL, David Ledbetter H, Kallen RG, Barchi RL (1991) Assignment of a human skeletal muscle sodium channel α-subunit gene (SCN4A) to 17q23.1–25.3. Genomics 9:555–556

Gohl DM, Silies MA, Gao XJ, Bhalerao S, Luongo FJ, Lin C-C, Potter CJ, Clandinin TR (2011) A versatile in vivo system for directed dissection of gene expression patterns. Nature Methods 8:231–237

Golding NL, Spruston N (1998) Dendritic Sodium Spikes Are Variable Triggers of Axonal Action Potentials in Hippocampal CA1 Pyramidal Neurons. Neuron 21:1189–1200

Gong Z (2004) Two Interdependent TRPV Channel Subunits, Inactive and Nanchung, Mediate Hearing in Drosophila. Journal of Neuroscience 24:9059–9066

Göpfert MC, Albert JT, Nadrowski B, Kamikouchi A (2006) Specification of auditory sensitivity by Drosophila TRP channels. Nature Neuroscience 9:999–1000 Available at: http://www.nature.com/articles/nn1735 [Accessed October 26, 2021].

Grueber WB, Graubard K, Truman JW (2001) Tiling of the body wall by multidendritic sensory neurons in Manduca sexta. The Journal of Comparative Neurology 440:271–283

Grueber WB, Jan LY, Jan YN (2003) Different Levels of the Homeodomain Protein Cut Regulate Distinct Dendrite Branching Patterns of Drosophila Multidendritic Neurons. Cell 112:805– 818

Hartenstein V (1988) Development of Drosophila larval sensory organs: Spatiotemporal pattern of sensory neurones, peripheral axonal pathways and sensilla differentiation. Development 102:869–886.

Hassan BA, Bermingham NA, He Y, Sun Y, Jan Y-N, Zoghbi HY, Bellen HJ (2000) atonal Regulates Neurite Arborization but Does Not Act as a Proneural Gene in the Drosophila Brain. Neuron 25:549–561

Hill KG (1983) The physiology of locust auditory receptors. Journal of Comparative Physiology A 152:483–493

Hofmann T, Chubanov V, Gudermann T, Montell C (2003) TRPM5 Is a Voltage-Modulated and Ca2+-Activated Monovalent Selective Cation Channel. Current Biology 13:1153–1158

Hong C, Ganetzky B (1994) Spatial and temporal expression patterns of two sodium channel genes in Drosophila. The Journal of Neuroscience 14:5160–5169

Hu Y, Flockhart I, Vinayagam A, Bergwitz C, Berger B, Perrimon N, Mohr SE (2011) An integrative approach to ortholog prediction for disease-focused and other functional studies. BMC Bioinformatics 12:357.

Huang CY-M, Rasband MN (2018) Axon initial segments: structure, function, and disease. Ann N Y Acad Sci 1420:46–61

Huang J, Zhou W, Watson AM, Jan Y-N, Hong Y (2008) Efficient Ends-Out Gene Targeting In Drosophila. Genetics 180:703–707

Husain N, Pellikka M, Hong H, Klimentova T, Choe KM, Clandinin T, Tepass U (2006) The Agrin/Perlecan-Related Protein Eyes Shut Is Essential for Epithelial Lumen Formation in the *Drosophila* Retina. Developmental Cell 11:483–93

Jarman AP, Grau Y, Jan LY, Jan YN (1993) atonal is a proneural gene that directs chordotonal organ formation in the Drosophila peripheral nervous system. Cell 73:1307–1321

Jegla T, Nguyen MM, Feng C, Goetschius DJ, Luna E, van Rossum DB, Kamel B, Pisupati A, Milner ES, Rolls MM (2016) Bilaterian Giant Ankyrins Have a Common Evolutionary Origin and Play a Conserved Role in Patterning the Axon Initial Segment Desplan C, ed. PLOS Genetics 12:e1006457

Kamikouchi A, Shimada T, Ito K (2006) Comprehensive classification of the auditory sensory projections in the brain of the fruit fly *Drosophila melanogaster*. The Journal of Comparative Neurology 499:317–356

Kavlie RG, Kernan MJ, Eberl DF (2010) Hearing in Drosophila Requires TilB, a Conserved Protein Associated With Ciliary Motility. Genetics 185:177–188

Kim J, Chung YD, Park D, Choi S, Shin DW, Soh H, Lee HW, Son W, Yim J, Park C-S, Kernan MJ, Kim C (2003) A TRPV family ion channel required for hearing in Drosophila. Nature 424:81–84

Larkin A et al. (2021) FlyBase: updates to the *Drosophila melanogaster* knowledge base. Nucleic Acids Research 49:D899–D907

Lee J, Moon S, Cha Y, Chung YD (2010) Drosophila TRPN(= NOMPC) Channel Localizes to the Distal End of Mechanosensory Cilia Gonzalez C, ed. PLoS ONE 5:e11012

Lehnert BP, Baker AE, Gaudry Q, Chiang A-S, Wilson RI (2013) Distinct Roles of TRP Channels in Auditory Transduction and Amplification in Drosophila. Neuron 77:115–128

Li B, Li S, Zheng H, Yan Z (2021) Nanchung and Inactive define pore properties of the native auditory transduction channel in Drosophila. Proceedings of the National academy of Science 118 (49) e2106459118

Li T, Bellen HJ, Groves AK (2018) Using Drosophila to study mechanisms of hereditary hearing loss. DMM Disease Models and Mechanisms 11

Li T, Giagtzoglou N, Eberl DF, Jaiswal SN, Cai T, Godt D, Groves AK, Bellen HJ (2016) The E3 ligase Ubr3 regulates Usher syndrome and MYH9 disorder proteins in the auditory organs of Drosophila and mammals. Elife 5:1–23

Liang X, Madrid J, Saleh HS, Howard J (2011) NOMPC, a member of the TRP channel family, localizes to the tubular body and distal cilium of Drosophila campaniform and chordotonal receptor cells. Cytoskeleton 68:1–7

Lin W-H, Gunay C, Marley R, Prinz AA, Baines RA (2012) Activity-Dependent Alternative Splicing Increases Persistent Sodium Current and Promotes Seizure. Journal of Neuroscience 32:7267–7277

Lin W-H, Wright DE, Muraro NI, Baines RA (2009) Alternative Splicing in the Voltage-Gated Sodium Channel DmNa _v_ Regulates Activation, Inactivation, and Persistent Current. Journal of Neurophysiology 102:1994–2006

Liu L, Yermolaieva O, Johnson WA, Abboud FM, Welsh MJ (2003) Identification and function of thermosensory neurons in Drosophila larvae. Nature Neuroscience 6:267–273

Lorincz A, Nusser Z (2010) Molecular Identity of Dendritic Voltage-Gated Sodium Channels. Science (1979) 328:906–909

Losonczy A, Magee JC (2006) Integrative Properties of Radial Oblique Dendrites in Hippocampal CA1 Pyramidal Neurons. Neuron 50:291–307

Luo L, Liao YJ, Jan LY, Jan YN (1994) Distinct morphogenetic functions of similar small GTPases: Drosophila Drac1 is involved in axonal outgrowth and myoblast fusion. Genes & Development 8:1787–1802

Mackenzie FE, Parker A, Parkinson NJ, Oliver PL, Brooker D, Underhill P, Lukashkina VA, Lukashkin AN, Holmes C, Brown SDM (2009) Analysis of the mouse mutant *Cloth-ears* shows a role for the voltage-gated sodium channel *Scn8a* in peripheral neural hearing loss. Genes, Brain and Behavior 8:699–713

Matta JA, Ahern GP (2007) Voltage is a partial activator of rat thermosensitive TRP channels. The Journal of Physiology 585:469–482

Montell C (2021) *Drosophila* sensory receptors—a set of molecular Swiss Army Knives Truman J, ed. Genetics 217:1–34

Montell C, Birnbaumer L, Flockerzi V, Bindels RJ, Bruford EA, Caterina MJ, Clapham DE, Harteneck C, Heller S, Julius D, Kojima I, Mori Y, Penner R, Prawitt D, Scharenberg AM, Schultz G, Shimizu N, Zhu MX (2002) A Unified Nomenclature for the Superfamily of TRP Cation Channels. Molecular Cell 9:229–231

Morris CE (2011) Voltage-Gated Channel Mechanosensitivity: Fact or Friction? Frontiers in Physiology 2:1–10

Morris CE, Juranka PF (2007) Nav Channel Mechanosensitivity: Activation and Inactivation Accelerate Reversibly with Stretch. Biophysical Journal 93:822–833

Nelson HB and Laughon A. (1993) Drosophila glial architechture and development: analysis using a collection of new cell specific markers. Roux’s Archives of Developmental Biology 202:341–354

Nilius B, Prenen J, Droogmans G, Voets T, Vennekens R, Freichel M, Wissenbach U, Flockerzi V (2003) Voltage Dependence of the Ca2+-activated Cation Channel TRPM4. Journal of Biological Chemistry 278:30813–30820

Ogiwara I, Ito K, Sawaishi Y, Osaka H, Mazaki E, Inoue I, Montal M, Hashikawa T, Shike T, Fujiwara T, Inoue Y, Kaneda M, Yamakawa K (2009) De novo mutations of voltage-gated sodium channel II gene SCN2A in intractable epilepsies. Neurology 73:1046–1053

Oldfield BP, Hill KG (1986) Functional organization of insect auditory sensilla. Journal of Comparative Physiology A 158:27–34

Orgogozo V, Grueber WB (2005) FlyPNS, a database of the Drosophila embryonic and larval peripheral nervous system. BMC Dev Biol 5:4

Owsianik G, Talavera K, Voets T, Nilius B (2006) Permeation and selectivity of TRP channels. Annual Review of Physiology 68:685–717

Pauron D, Barhanin J, Lazdunski M (1985) The voltage-dependent Na+ channel of insect nervous system identified by receptor sites for tetrodotoxin, and scorpion and sea anemone toxins. Biochemical and Biophysical Research Communications 131:1226–1233

Pfeiffer BD, Ngo T-TB, Hibbard KL, Murphy C, Jenett A, Truman JW, Rubin GM (2010) Refinement of Tools for Targeted Gene Expression in Drosophila. Genetics 186:735–755

Ravenscroft TA, Janssens J, Lee P-T, Tepe B, Marcogliese PC, Makhzami S, Holmes TC, Aerts S, Bellen HJ (2020) *Drosophila* Voltage-Gated Sodium Channels Are Only Expressed in Active Neurons and Are Localized to Distal Axonal Initial Segment-like Domains. The Journal of Neuroscience 40:7999–8024

Rothwell WF, Sullivan W (2007) Fixation of Drosophila Embryos. Cold Spring Harbor Protocols 2007:pdb.prot4827

Roy M, Sivan-Loukianova E, Eberl DF (2013) Cell-type-specific roles of Na+/K+ ATPase subunits in Drosophila auditory mechanosensation. Proceedings of the National Academy of Sciences 110:181–186

Schneider CA, Rasband WS, Eliceiri KW (2012) NIH Image to ImageJ: 25 years of image analysis. Nature Methods 9:671–675

Schrader Š, Merritt DJ (2007) Dorsal longitudinal stretch receptor of Drosophila melanogaster larva – Fine structure and maturation. Arthropod Structure & Development 36:157–169

Shcherbatko A, Ono F, Mandel G, Brehm P (1999) Voltage-Dependent Sodium Channel Function Is Regulated Through Membrane Mechanics. Biophysical Journal 77:1945–1959

Shearin HK, Macdonald IS, Spector LP, Stowers RS (2014) Hexameric GFP and mCherry Reporters for the *Drosophila* GAL4, Q, and LexA Transcription Systems. Genetics 196:951–960

Singhania A, Grueber WB (2014) Development of the embryonic and larval peripheral nervous system of *Drosophila*. Wiley Interdisciplinary Reviews: Developmental Biology 3:193–210

Stocker RF (1994) The organization of the chemosensory system in Drosophila melanogaster: a rewiew. Cell and Tissue Research 275:3–26

Strege PR, Mercado-Perez A, Mazzone A, Saito YA, Bernard CE, Farrugia G, Beyder A (2019) SCN5A mutation G615E results in NaV1.5 voltage-gated sodium channels with normal voltage-dependent function yet loss of mechanosensitivity. Channels (Austin) 13:287–298

Stuart G, Schiller J, Sakmann B (1997) Action potential initiation and propagation in rat neocortical pyramidal neurons. The Journal of Physiology 505:617–632

Sun Q, Srinivas K v, Sotayo A, Siegelbaum SA (2014) Dendritic Na+ spikes enable cortical input to drive action potential output from hippocampal CA2 pyramidal neurons. Elife 3:1–24

Suslak TJ, Armstrong JD, Jarman AP (2011) A general mathematical model of transduction events in mechano-sensory stretch receptors. Network: Computation in Neural Systems 22:133– 142

Suslak TJ, Watson S, Thompson KJ, Shenton FC, Bewick GS, Armstrong JD, Jarman AP (2015) Piezo Is Essential for Amiloride-Sensitive Stretch-Activated Mechanotransduction in Larval Drosophila Dorsal Bipolar Dendritic Sensory Neurons McCabe BD, ed. PLOS ONE 10:e0130969

Suzuki DT, Grigliatti T, Williamson R (1971) Temperature-Sensitive Mutations in Drosophila melanogaster, VII. A Mutation (parats) Causing Reversible Adult Paralysis. Proceedings of the National Academy of Sciences 68:890–893

Swerup C, Rydqvist B (1996) A mathematical model of the crustacean stretch receptor neuron. Biomechanics of the receptor muscle, mechanosensitive ion channels, and macrotransducer properties. Journal of Neurophysiology 76:2211–2220

Tabarean I v., Juranka P, Morris CE (1999) Membrane Stretch Affects Gating Modes of a Skeletal Muscle Sodium Channel. Biophysical Journal 77:758–774

Thevenaz P, Ruttimann UE, Unser M (1998) A pyramid approach to subpixel registration based on intensity. IEEE Transactions on Image Processing 7:27–41

Todi S v, Sharma Y, Eberl DF (2004) Anatomical and molecular design of the Drosophila antenna as a flagellar auditory organ. Microsc Res Tech 63:388–399

Tracey WD, Wilson RI, Laurent G, Benzer S (2003) painless, a Drosophila Gene Essential for Nociception. Cell 113:261–273

Trudeau MM (2006) Heterozygosity for a protein truncation mutation of sodium channel SCN8A in a patient with cerebellar atrophy, ataxia, and mental retardation. Journal of Medical Genetics 43:527–530

Tsubouchi A, Caldwell JC, Tracey WD (2012) Dendritic Filopodia, Ripped Pocket, NOMPC, and NMDARs Contribute to the Sense of Touch in Drosophila Larvae. Current Biology 22:2124– 2134

Veeramah KR, O’Brien JE, Meisler MH, Cheng X, Dib-Hajj SD, Waxman SG, Talwar D, Girirajan S, Eichler EE, Restifo LL, Erickson RP, Hammer MF (2012) De Novo Pathogenic SCN8A Mutation Identified by Whole-Genome Sequencing of a Family Quartet Affected by Infantile Epileptic Encephalopathy and SUDEP. The American Journal of Human Genetics 90:502–510

Venken KJT, Schulze KL, Haelterman NA, Pan H, He Y, Evans-Holm M, Carlson JW, Levis RW, Spradling AC, Hoskins RA, Bellen HJ (2011) MiMIC: a highly versatile transposon insertion resource for engineering Drosophila melanogaster genes. Nature Methods 8:737–743

Vervoort M, Merritt DJ, Ghysen A, Dambly-Chaudiere C (1997) Genetic basis of the formation and identity of type I and type II neurons in Drosophila embryos. Development 124:2819– 2828

Voets T, Droogmans G, Wissenbach U, Janssens A, Flockerzi V, Nilius B (2004) The principle of temperature-dependent gating in cold- and heat-sensitive TRP channels. Nature 430:748– 754

Wang H, Foquet B, Dewell RB, Song H, Dierick HA, Gabbiani F (2020) Molecular characterization and distribution of the voltage-gated sodium channel, Para, in the brain of the grasshopper and vinegar fly. Journal of Comparative Physiology A 206:289–307

Wang X, Zhang MW, Kim JH, Macara AM, Sterne G, Yang T, Ye B (2015) The Kruppel-Like Factor Dar1 Determines Multipolar Neuron Morphology. Journal of Neuroscience 35:14251– 14259

Warren B, Matheson T (2018) The Role of the Mechanotransduction Ion Channel Candidate Nanchung-Inactive in Auditory Transduction in an Insect Ear. The Journal of Neuroscience 38:3741–3752

Wei Z, Lin B-J, Chen T-W, Daie K, Svoboda K, Druckmann S (2020) A comparison of neuronal population dynamics measured with calcium imaging and electrophysiology Gutkin BS, ed. PLOS Computational Biology 16:e1008198

Zhang W, Yan Z, Jan LY, Jan YN (2013) Sound response mediated by the TRP channels NOMPC, NANCHUNG, and INACTIVE in chordotonal organs of Drosophila larvae. Proceedings of the National Academy of Sciences 110:13612–13617

Zheng J (2013) Molecular Mechanism of TRP Channels. In: Comprehensive Physiology, pp 221–242. Wiley.

Zhong L, Hwang RY, Tracey WD (2010) Pickpocket Is a DEG/ENaC Protein Required for Mechanical Nociception in Drosophila Larvae. Current Biology 20:429–434

